# Three points of consideration before testing the effect of patch connectivity on local species richness: patch delineation, scaling and variability of metrics

**DOI:** 10.1101/640995

**Authors:** Fabien Laroche, Manon Balbi, Théophile Grébert, Franck Jabot, Frédéric Archaux

## Abstract

The Theory of Island Biogeography (TIB) promoted the idea that species richness within sites depends on site connectivity, i.e. its connection with surrounding potential sources of immigrants. TIB has been extended to a wide array of fragmented ecosystems, beyond archipelagoes, surfing on the analogy between habitat patches and islands and on the patch-matrix framework. However, patch connectivity often little contributes to explaining species richness in empirical studies. Before interpreting this trend as questioning the broad applicability of TIB principles, one first needs a clear identification of methods and contexts where strong effects of patch structural connectivity are likely to occur. Here, we use spatially explicit simulations of neutral metacommunities to show that patch connectivity effect on local species richness is maximized under a set of specific conditions: (i) patch delineation should be fine enough to ensure that no dispersal limitation occurs within patches, (ii) patch connectivity indices should be scaled according to target organisms’ dispersal distance and (iii) the habitat amount around sampled sites (within a distance adapted to organisms’ dispersal) should be highly variable. When those three criteria are met, the absence of an effect of connectivity on species richness should be interpreted as contradicting TIB hypotheses

## Introduction

Since the Theory of Island Biogeography (TIB) [1], it is commonly acknowledged that species presence within local communities depends on their ability to immigrate, and that geographic isolation of communities can negatively affect species richness. Initially devised for insular ecosystems, TIB principles have been extended to terrestrial ecosystems (see [2,3] for reviews and critical appraisal). This led to studying how the availability of suitable habitat nearby, called “patch structural connectivity” [4], can act as a source of immigrants and affect species richness within local communities. Extending TIB relied on adopting a “patch-matrix” description of habitat in space, where one decomposes the map of some suitable habitat into patches that correspond to potential communities (analogous to islands in an archipelago), the rest of space being considered unhospitable for the species under study

Most of the tests regarding the species diversity patterns predicted by the TIB have focused on the shape of the species richness – patch area curve, and studied how patch connectivity can modulate this relationship [5]. There is unfortunately no systematic review or meta-analysis about the species richness – patch connectivity relationship *per se*. Scattered empirical studies from the literature suggest that connectivity effects on species richness are variable in the field. The direction and strength of the relationship seems to depend, among other factors, on the dispersal distance [6,7], the trophic level [8–10] and the degree of generalism [11,12] of considered species groups, as well as on the perturbation history of sites [13]

Clearer syntheses are available when considering the effect of patch connectivity on individual species presences rather than species richness in itself. In a meta-analysis of 1,015 empirical studies within terrestrial systems, Prugh et al. [14] evidenced that patch structural connectivity, measured as distance to the nearest patch, tends to have weak predictive power on species presence within patches (median deviance explained equaled about 20%). Another review of 122 empirical studies [15], which covered terrestrial and aquatic systems and analyzed the presence or abundance of 954 species, evidenced that the effects of local environmental conditions within a patch on species presence or abundance occurred more frequently (71% of species analyses) than the effects of patch structural connectivity (55% of species analyses)

Former studies thus tend to suggest that patch structural connectivity seems to be a non-robust and relatively weak predictor of species richness, which potentially questions the role of immigration as an important process in community assembly within fragmented habitats. According to Prugh et al. [14], the limited success of patch connectivity indices may come from several conceptual flaws of applying the TIB framework to non-archipelago landscapes: (i) inadequately using structural connectivity indices based on surrounding habitat rather than functional connectivity indices based on surrounding populations (i.e. habitat being actually occupied by target species); (ii) inadequately delineating patches for species harboring multiple life stages with contrasted requirements (e.g., juveniles living in aquatic habitats and adults living in terrestrial habitats); (iii) overlooking the type of matrix surrounding the habitat patch, hence questioning the validity of the patch-matrix framework for terrestrial systems. In the same vein, Cook et al. [16] also suggested that an important fraction of species found in patches could also survive and even thrive in the matrix, hence explaining the failure of TIB applications to non-archipelago landscapes

However, the limited success of patch structural connectivity indices may also stem from several methodological limits. Thornton et al. [15] mentioned for instance the problem of using inadequate patch structural connectivity metrics, emphasizing that buffer indices are generally more performant than widely used isolation metrics (a point that was also raised by several other studies [17,18]). Second, inadequate scaling of indices with respect to target organisms dispersal distance may also drive down the explanatory power of patch connectivity indices upon species presence/absence, community diversity and presumably species richness. The higher the dispersal distance of species, the larger the scaling of indices should be to reach the best possible explanatory power, as demonstrated by several simulation studies [19,20]. Third, patches are often built through lumping together sets of contiguous habitat pixels on a land cover map, following a “vector map” perspective [21]. However, this approach brings no guarantee that the resulting spatial entities have the appropriate size to constitute potential communities for target organisms, and considering entities with inadequate spatial resolution with respect to target processes is known to erode expected patterns [22]. Fourth, empirical studies about connectivity effects may have suffered from a lack of statistical power due to low sample size (i.e. low number of patches) [15]

Interpreting the limited predictive power of patch structural connectivity as an explanatory failure of the TIB framework is valid only when methods used and landscape context should theoretically foster large effect sizes. Therefore, before questioning the first principles of the TIB framework, one first needs identifying which methods for measuring patch structural connectivity and which properties of the habitat spatial distribution of studied systems should yield strong effects of patch structural connectivity on local species richness

In our analysis, we focused on how the patch delineation, the type of patch connectivity index, the scaling of indices with species dispersal distance, and the variability of indices within landscapes affect the explanatory power of patch structural connectivity on local species richness. We aimed at deriving good practices with respect to these four points. In particular, we made three hypotheses. First, we expected that delineating habitat patches with sizes matching the typical extension of communities should allow higher explanatory power of patch connectivity indices than a coarser patch delineation. This stems from the fact that delineating patches larger than communities implies that a part of the species dispersal limitation effects on local species richness occurs within patches and is thus invisible to analyses focusing on connectivity among patches. Second, we expected species with larger dispersal distance to require patch connectivity indices accounting for habitat patches located further away from the focal patch (i.e. larger scaling) in order to maximize connectivity explanatory power. Third and last, we expected a strong explanatory power of connectivity indices to arise only if connectivity indices are highly variable among patches. Otherwise, the effect of connectivity on species richness is likely to be diluted in other independent sources of variation of species richness among sites, thus becoming impossible to detect in studies with limited statistical power

We used a virtual ecologist approach [23] relying on metacommunity simulations in a spatially-explicit model to test these hypotheses. Virtual datasets stemming from such models constitute an ideal context to assess the impact of our factors of interest, since they offer perfect control of the spatial distribution of habitat and the ecological features of species. In particular, we only included processes related to the TIB (immigration, ecological drift; [24]), thus maximizing our ability to study how methodological choices and landscape features affect the explanatory power of patch structural connectivity. We anticipated that explanatory powers generated by this approach would necessarily be an over-estimation of what occurs in real ecosystems, where many processes unrelated to TIB may also be at work. This implies in particular that settings negatively affecting the explanatory power of patch structural connectivity in our simulation study have very little chance to yield strong explanatory power in empirical studies, and cannot be used to criticize the hypotheses of TIB

## Methods

### Landscape generation

We considered binary landscapes made up of suitable habitat cells and inhospitable matrix cells. We generated virtual landscapes composed of 100×100 cells using a midpoint-displacement algorithm [25] which allowed us covering different levels of habitat quantity and fragmentation. The proportion of habitat cells varied according to three modalities (10%, 20%, and 40% of the landscapes). The spatial aggregation of habitat cells varied independently, and was controlled by the Hurst exponent (0.1, 0.5, and 0.9 in increasing order of aggregation; see Fig. S1 for examples). We generated ten replicates for each of these nine landscape types, resulting in 90 landscapes. Higher values of the Hurst exponent for a given value of habitat proportion increased the size of contiguous habitat patches and decreased the number of distinct patches (Fig. S2). Higher habitat proportion for a constant Hurst exponent value also resulted in higher mean size of contiguous habitat patches

### Neutral metacommunity simulations

We simulated spatially explicit neutral metacommunities on the virtual heterogeneous landscapes described above. We resorted to using a spatially explicit neutral model of metacommunities, where all species had the same dispersal distance. We used a discrete-time model where the metacommunity changed by steps. All habitat cells were occupied, and community dynamics in each habitat cell followed a zero-sum game, so that habitat cells always harbored 100 individuals at the beginning of a step. One step was made up of two consecutive events. Event 1: 10% of individuals diex in each cell – they were picked at random. Event 2: dead individuals were replaced by the same number of recruited individuals that were randomly drawn from a multinomial distribution, each species having a weight equal to 0.01×χ_i_ + ∑_k_ A_ik_ exp(-d_kf_ /λ_s_) where χ_i_ is the relative abundance of species i in the regional pool, A_ik_ is the local abundance of species i in habitat cell k, d_kf_ is the distance (in cell unit) between the focal habitat cell f and the source habitat cell k, λ_s_ is a parameter defining species dispersal distances and the sum is over all habitat cells k of the landscape

The regional pool was an infinite pool of migrants representing biodiversity at larger spatial scales than the focal landscape, it contained 100 species, the relative abundances of which (χ_i_s) were sampled once for all at the beginning of the simulation in a Dirichlet distribution with concentration parameters α_i_ equal to 1 (with i from 1 to 100)

Distances between habitat pixels (d_kf_) were defined as the Euclidean distance on a torus, to remove unwanted border effects in metacommunity dynamics. Metacommunities were simulated with three levels of species dispersal distance: λ_s_ = 0.25, 0.5, 1 cell, which corresponded to median dispersal distance of 0.6, 0.7, 0.9 cell and average dispersal distance of 0.6, 0.8, 1.2 cells. The 95% quantile of dispersal distance corresponded to 1.2, 1.7, 3.1 cells respectively

For a given landscape replicate, metacommunity replicates were obtained by recording the state of a metacommunity at various dates in one forward in time simulation, with 1000 burn-in steps and 500 steps between each replicate. The recorded state of the metacommunity included the abundances of each species in each habitat cell. We performed 10 replicates for each dispersal distance value and in each simulated landscape. In total, we obtained 2,700 metacommunity replicates (3 Hurst exponent values × 3 habitat proportions × 3 species dispersal distances × 10 landscape replicates × 10 metacommunity replicates)

### Patch connectivity indices

For each habitat cell of the 90 simulated landscapes, we computed three types of patch connectivity indices (Table 1; Fig. 1A): *Buffer*, *dF* and *dIICflux*. *Buffer* indices corresponded to the proportion of area covered by habitat within circles of different radius (r_buf_ = 1, 2, 4, 5, 8 cells) around the focal cell

**Table 1.** Patch connectivity indices considered in the study

| Index | Definition | Ref. |
| --- | --- | --- |
| *Buffer* | $\text{buf}_k = \frac{a}{\pi r^2} \sum_{i=1, i \neq k}^n 1_{d_{ik} \leq r}$ | [17] |
| *dIICflux* | $dIICflux_k = \frac{100}{IIC} \left[ 2 \sum_{i=1, i \neq k}^n \frac{a_k a_i}{1 + nl_{ik}} \right]$ | [28] |
| *dF* | $dF_k = 2 \sum_{i=1, i \neq k}^n w_{ik}$ | [21,26] |

Notations — $n$ : total number of nodes (patches or cells) in a graph; $a$ : area of a cell; $a_i$ : area of patch $i$ ; $r$ : radius of *Buffer*; $nl_{ij}$ : shortest path between nodes $i$ and $j$ in a binary graph; $IIC = \sum_{i=1}^n \sum_{j=1}^n a_i a_j / (1 + nl_{ij})$ : integral index of connectivity of a graph; $d_{ij}$ : Euclidean distance between nodes $i$ and $j$ ; $w_{ij}$ : probability weight of the link between nodes $i$ and $j$ in a weighted graph.

**Figure 1.**
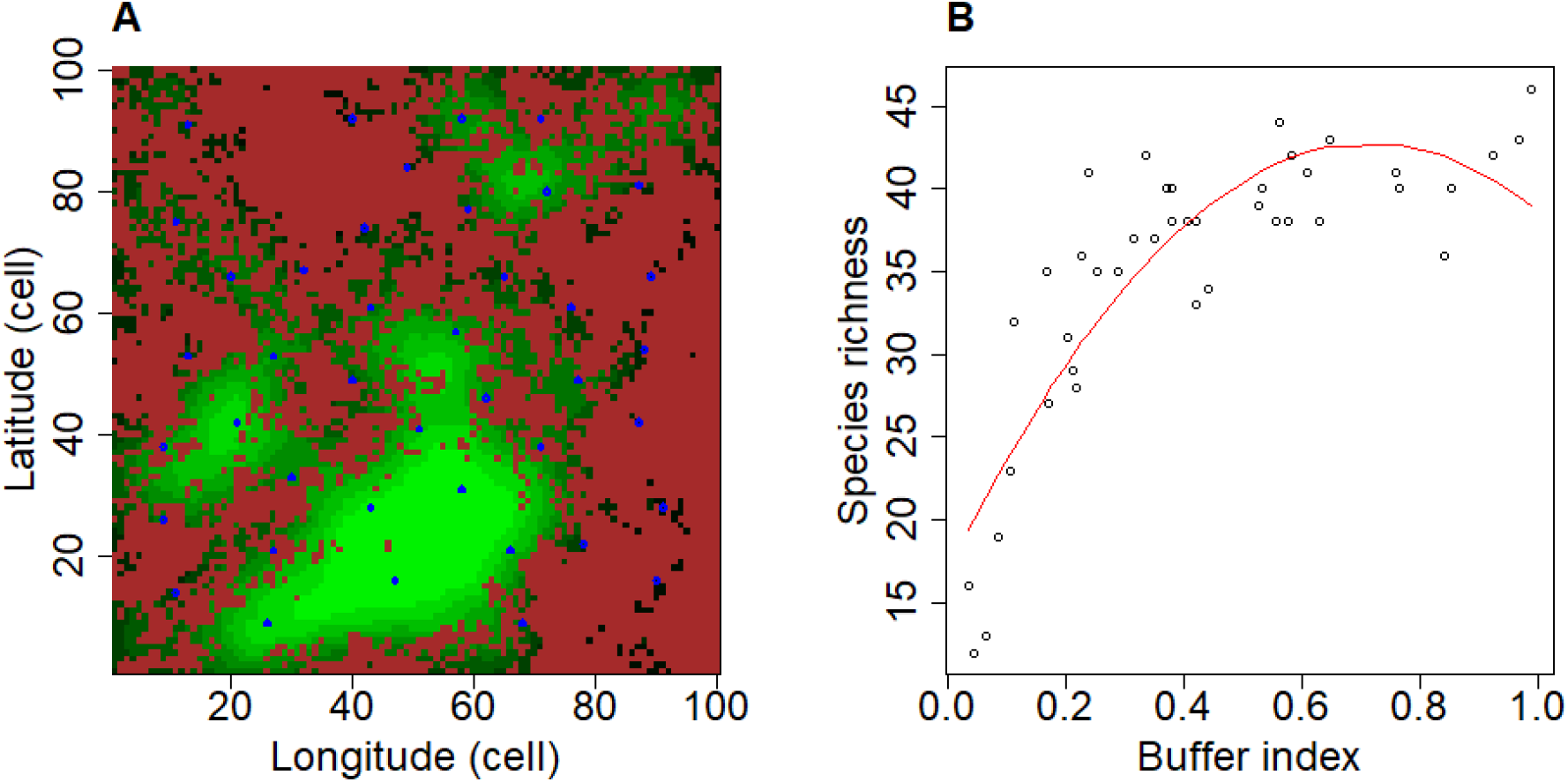
Example of analysis of the explanatory power of Buffer index in a virtual dataset. Panel A: a virtual landscape obtained through midpoint displacement algorithm, with controlled habitat proportion (here 0.4) and Hurst exponent (here 0.1). Brown cells stands for unhospitable matrix. Green cells denote habitat cells. Lighter cells harbor a higher patch connectivity *Buffer* index (here computed with radius 8 cells). Blue circles show sampled cells. Panel B: relationship between *Buffer* index and species richness in sampled cells for a metacommunity replicate within the landscape of panel A. The relationship was analyzed using a quadratic model (red curve), and the R^2^ of the model, *R2_spec_,* was recorded for future analyses. The species dispersal distance used to simulate the metacommunity replicate presented here was λ_s_ = 1 cell

The computation of *dIICflux* and *dF* relied on delineating habitat “patches”. We alternatively considered two delineation approaches (Fig. S3): patches were defined either as single habitat cells in the landscape (“fine” patch delineation) or as groups of contiguous habitat cells (“coarse” patch delineation). With fine patch delineation, each patch corresponded to a single community. With coarse patch delineation, patches contained as many communities as cells forming the patches. For each approach, pairs of patches obtained were then connected by links. Links’ weights w_ij_ between nodes i and j in the network decreased according to the formula exp(-d_ij_/λ_c_), where d_ij_ is the Euclidean distances between nodes i and j and λ_c_ is a scale parameter [21,26]. λ_c_ may be interpreted as the hypothesized scale of dispersal distance of target organisms in the landscape (which may differ from the “true” simulated scale of dispersal distance, which is λ_s_). We considered four scale parameter values (λ_c_ = 0.25, 0.5, 1 and 2 cells). *dF* quantified the sum of edges weights between the focal patch and all the other patches. *dIICflux* considered a binary graph, where each node pair was considered either connected (1) or not (0) relatively to a minimal link weight w_min_ = 0.005. Scale parameters λ_c_ = 0.25, 0.5, 1 and 2 cells thus lead to connect all pairs of habitat cells separated by a distance inferior to 1.3, 2.6, 5.3 and 10.6 cells respectively. *dIICflux* captured a notion of node centrality, like *dF*, but based on topological distance in the graph rather than Euclidean distance. All indices were computed with Conefor 2.7 (command line version for Linux, furnished by S. Saura, soon publicly available on www.conefor.org; [27])

Altogether, in each habitat cell of each simulated landscape, we computed 5 *Buffer* indices corresponding to distinct radii, 8 *dF* indices corresponding to 4 scale parameters at both fine and coarse patch delineation, and 8 *dIICflux* indices also corresponding to 4 scale parameters at both fine and coarse patch delineation. In total, this led to 21 patch connectivity indices per habitat cell

### Sampling design

For each simulated landscape, we defined a set of sampled cells, including habitat cells away from each other’s for a minimal distance of 12 cells, to reduce spatial auto-correlation (e.g. Fig. 1A). We also reduced potential landscape border effect by excluding cells near landscape borders (to a distance inferior or equal to eight cells, equivalent to the longest radius used for *Buffer* index, see below). Each landscape counted on average 25 sampled cells (CI-95% = [23, 27])

### General approach

For each of the 2,700 metacommunity recorded states, we computed species richness within habitat cells belonging to the sampling design. We thus obtained 21 × 2,700 = 56,700 relationships between a connectivity index and species richness. For each relationship, we computed the proportion of species richness variance explained by a quadratic function of the connectivity index (e.g. Fig. 1B). This proportion was obtained as the R^2^ coefficient of the linear model *Species richness ~ Patch connectivity + (Patch connectivity)_2_*. We henceforth refer to this coefficient as the “explanatory power” of the connectivity index, denoted “*R2_spec_*”. When considering coarse patch delineation, we additionally included the effect of the area of the patch in the explanatory power of connectivity, meaning that *R2*_*spec*_ was obtained as the R^2^ coefficient of the model *Species richness ~ Patch connectivity + (Patch connectivity)^2^ + Patch area + (Patch area)^2^.* We did so to ensure fair comparison with the fine patch delineation

We applied linear models on *R2*_*spec*_ values to test our three hypotheses, as detailed below

### Testing hypothesis 1: patch delineation effect

Our first hypothesis was that the explanatory power of connectivity indices should be optimal when patch delineation matches the geographical scale of studied communities, hence ensuring no within-patch dispersal limitation. In our simulations, the size of habitat cells within a land cover map matches the community size of species, therefore we predicted that lumping contiguous habitat cells together into larger patches would deteriorate the explanatory power of indices with respect to species richness. We tested this hypothesis by exploring the effect of patch delineation on *R2*_*spec*_ for *dF* and *dIICflux* indices. *Buffer* indices were not considered in this analysis because they did not depend on patch delineation

In each of the 2,700 simulated datasets, we recorded *R2*_*spec*_ for *dF* or *dIICflux* computed with a fine and coarse patch delineation. We controlled for the variation of *R2*_*spec*_ due to index scaling (which are analyzed separately in the next section) by focusing, for each index and each patch delineation strategy, on the highest value out of the four distinct *R2*_*spec*_ values in the present analysis. We thus obtained 2,700 datasets × 2 patch delineation × 2 indices = 10,800 *R2*_*spec*_ values

We separately analyzed the *R2*_*spec*_ values of *dF* and *dIICflux*. We expected *R2*_*spec*_ to be significantly higher at fine patch delineation, which we tested using the model *R2_spec_ ~ patch delineation*. We also expected the positive effect of switching from coarse to fine patch delineation to increase when Hurst exponent (i.e. habitat aggregation) or habitat proportion increase, because patches (sets of contiguous cells here) become larger on average, leading to stronger dispersal limitation effects within patches. We tested this second hypothesis using linear models with interactions: *R2_spec_ ~ patch delineation* × *Hurst exponent* and *R2_spec_ ~ patch delineation* × *habitat proportion.* At last, we expected the positive effect of switching from coarse to fine patch delineation to decrease when species dispersal distance increases, because dispersal limitation within patches weakens. We tested this last hypothesis using the model: *R2_spec_~patch delineation* × *dispersal.* In the three latter linear models, the interacting covariate (Hurst exponent, habitat proportion and species dispersal respectively) was considered as a factor

### Testing hypothesis 2: index scaling effect

Our second hypothesis was that the scaling of patch connectivity indices maximizing the explanatory power upon species richness should increase with dispersal distance of target organisms. We tested it by recording, in the 2,700 simulated datasets, *R2*_*spec*_ for *Buffer*, *dIICflux* and *dF* patch connectivity indices computed with a fine patch delineation at each scaling parameter value. We thus obtained 2,700 datasets × 3 indices × 4 or 5 scaling parameter values [4 for *dF* and *dIICflux*, 5 for *Buffer*] = 35,100 *R2*_*spec*_ values. We then built one linear model per index type (*Buffer*, *dF* or *dIICflux*), where *R2*_*spec*_ was the dependent variable, modelled as a function of species dispersal distance in interaction with index scale parameter *R2_spec_ ~ dispersal* × *scaling value*. For each species dispersal distance, we could then identify the scaling of indices maximizing *R2_spec_*, which we call the “optimal” scaling. We expected the optimal scaling of indices to increase with the dispersal distance of species, following previously published results in the literature [19,20]

### Testing hypothesis 3: connectivity variability effect

Our third and last hypothesis was that a higher variability of patch connectivity indices among sampled sites should increase the explanatory power of connectivity metrics upon species. We tested it by recording, in the 2,700 simulated datasets, the maximal value of *R2*_*spec*_ across scaling parameter values for *Buffer*, *dIICflux* and *dF* patch connectivity indices computed with a fine patch delineation. This generated 2,700 virtual datasets × 3 index types = 8,100 *R2*_*spec*_ values. Then we explored separately for each index at each species dispersal distance how the coefficient of variation of patch connectivity indices affected *R2_spec_*. For each index, we computed the average value of *R2*_*spec*_ with optimal scaling across the 10 metacommunity replicates associated to one landscape and one dispersal distance level (*avR2*_*spec*_ below). We computed the corresponding average coefficient of variation of the patch connectivity index with optimal scaling (*avCV*). Thus, we obtained 3 Hurst exponent × 3 habitat proportion × 3 dispersal distance × 10 landscape replicates = 270 pairs of *avCV* and *avR2*_*spec*_ values. We analyzed the relationship between these quantities using the linear model *logit(avR2_spec_) ~ log(avCV)*, where we applied log(.) and logit(.) transformations to dependent and independent variables respectively to obtain linearity and homoscedasticity of the relationship. We expected landscapes with higher *avCV* to yield higher *avR2_spec_*

We additionally tested whether the effects of habitat aggregation and habitat proportion on *R2*_*spec*_ were completely mediated by the coefficient of variation of the connectivity index. To do so, we added the interaction of habitat aggregation and habitat proportion in the above-mentioned linear model (i.e. considering *logit(avR2_spec_)~log(avCV)+ Hurst exponent × habitat proportion*). A significant improvement of the model fit would have suggested that habitat aggregation and habitat proportion modulated the explanatory power of connectivity indices beyond their effect on its coefficient of variation among sampled sites

## Results

The median of the 56,700 *R2*_*spec*_ values obtained from our simulations was 0.65, suggesting that the explanatory power of patch connectivity indices was generally strong. However, the explanatory power fluctuated a lot around the median value: 2.5% of *R2*_*spec*_ values were below 0.07 while another 2.5% were above 0.94

### Hypothesis 1: patch delineation effect

For both *dF* and *dIICflux*, using a fine patch delineation yielded higher *R2*_*spec*_ than using a coarse patch delineation (*dF*: +0.19 on average, s.e.=0.005, *p*<2e-16; *dIICflux*: +0.08 on average, s.e.=0.006, *p*<2e-16)

For *dF* index, high Hurst exponent (high habitat aggregation) significantly increased the positive effect of refining patch delineation on *R2*_*spec*_ compared to medium or low Hurst exponent (*F*-test; *F*=3.8, *p*=0.02). However, this modulation had a limited effect size: for high Hurst exponent, *R2*_*spec*_ increased by +0.21 (s.e.=0.009) with fine delineation, while it increased by +0.18 (s.e.=0.009) for medium or low Hurst exponent. A larger proportion of habitat in the landscape significantly increased the positive effect of refining patch delineation on *R2*_*spec*_ (*F*-test; *F*=16.6, *p*=6e-8): the effect of refining patch delineation on *R2*_*spec*_ reached +0.23 (s.e.=0.009) for a habitat proportion of 0.4 while it equaled +0.16 (s.e.=0.009) only for a habitat proportion of 0.1 (Fig. 2A). Higher species dispersal distance decreased the positive effect of refining patch delineation on *R2*_*spec*_ (*F*-test; *F*=192, *p*<2e-16): the effect of refining patch delineation on *R2*_*spec*_ reached +0.28 (s.e.=0.008) when species had low dispersal distance while it equaled +0.07 (s.e.=0.008) when species had high dispersal distance (Fig. 2B)

**Figure 2.**
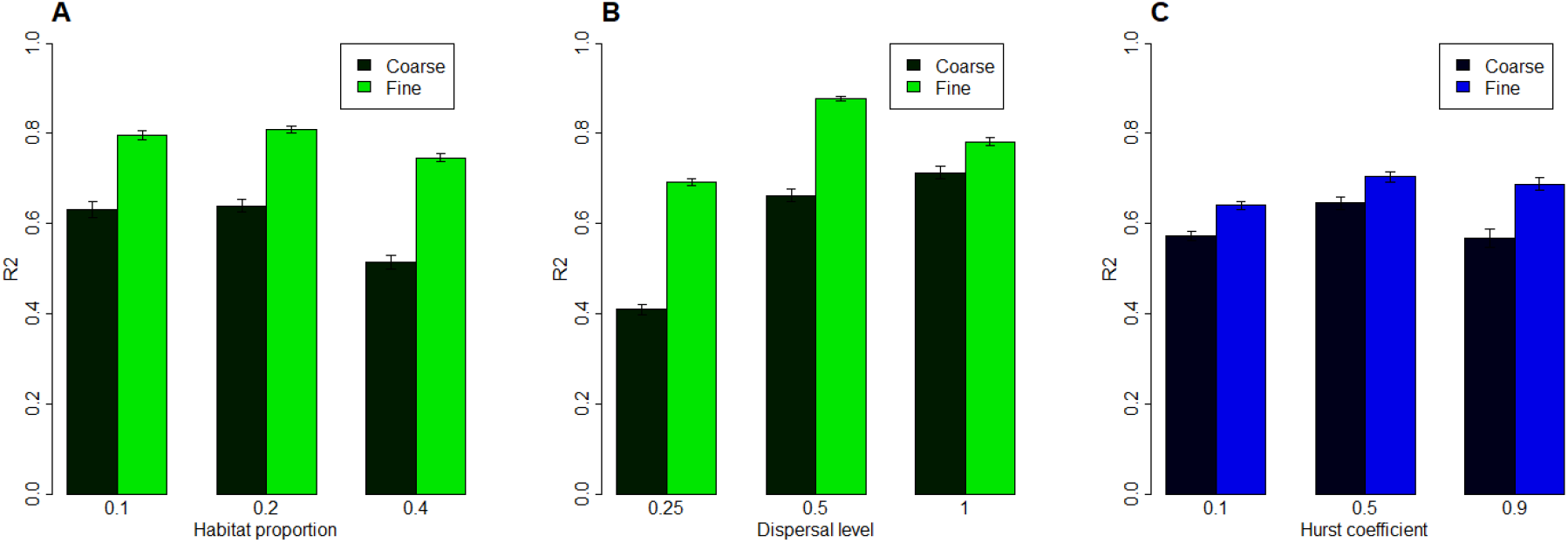
Hurst exponent, habitat proportion and species dispersal distance modulating the effect of refining patch delineation on the explanatory power of patch connectivity indices. Bars show the average *R2*_*spec*_ over simulated datasets for distinct levels of habitat proportion (panel A), species dispersal distance (panel B) and Hurst exponent (panel C), with asymptotic 95% confidence intervals (half width = 1.96 x standard error). Panel A and B come from the analysis of the *dF* index while Panel C comes from the analysis of *dIICflux*, hence the different color.

For *dIICflux* index, a higher Hurst exponent increased the positive effect of refining patch delineation on *R2*_*spec*_ (*F*-test; *F*=11.5, *p*=9e-6): the effect of refining patch delineation equaled +0.12 (s.e.=0.01) in highly aggregated landscapes with a Hurst exponent of 0.9. By contrast, the effect of refining patch delineation equaled +0.07 only (s.e.=0.01) in landscapes with a Hurst exponent of 0.1 (Fig. 2C) and +0.06 (s.e.=0.01) in landscapes with an intermediary Hurst exponent of 0.5. Habitat proportion and species dispersal distance did not significantly affect the effect of refining patch delineation on *R2_spec_*

### Hypothesis 2: index scaling effect

For *Buffer*, *dF* and *dIICflux*, the scaling parameter value yielding the highest *R2*_*spec*_ increased with species dispersal distance (Fig. 3). Because of our high number of simulations, the mean *R2*_*spec*_ obtained with optimal scaling was always significantly higher than mean values obtained with other scaling values. However, mean *R2*_*spec*_ rarely departed from the optimal scaling performance by more than one standard deviation, and it only happened for scaling parameter values very different from the optimal scaling (Fig. 3). Therefore, the magnitude of the variation of mean *R2*_*spec*_ between scaling parameter values could be considered as small compared to the intrinsic variability of *R2*_*spec*_ for a given scaling parameter value. The range of scaling parameters explored was not sufficient to obtain precise quantitative relationships between species dispersal distance and index scaling. For *Buffer* and *dF* indices, the optimal value could reach the higher end of the explored range for medium or high species dispersal distance, suggesting that the true optimal scaling value may actually be higher than the explored range. For *dIICflux* index, the optimal scaling value laid at the lower end of the explored range for low and medium species dispersal distances, suggesting that the true optimal scaling may actually be lower than the explored range. However, these results were sufficient to reveal that the relationship between species dispersal distance and optimal scaling is variable among the three types of index tested. In particular, the optimal scaling of *Buffer* radius (*r_buf_*) corresponded to about 8 times the true scale of species dispersal distance (*λ_s_*; Fig. 3A). The optimal scaling of *dF* indices (*λ_c_*) seemed to lie between 2 and 4 times the true scale of species dispersal distance (Fig. 3B), while the optimal scaling parameter of *dIICflux* indices (*λ_c_*) seemed to be about 0.5 times the true scale of species dispersal distance (Fig. 3C)

**Figure 3.**
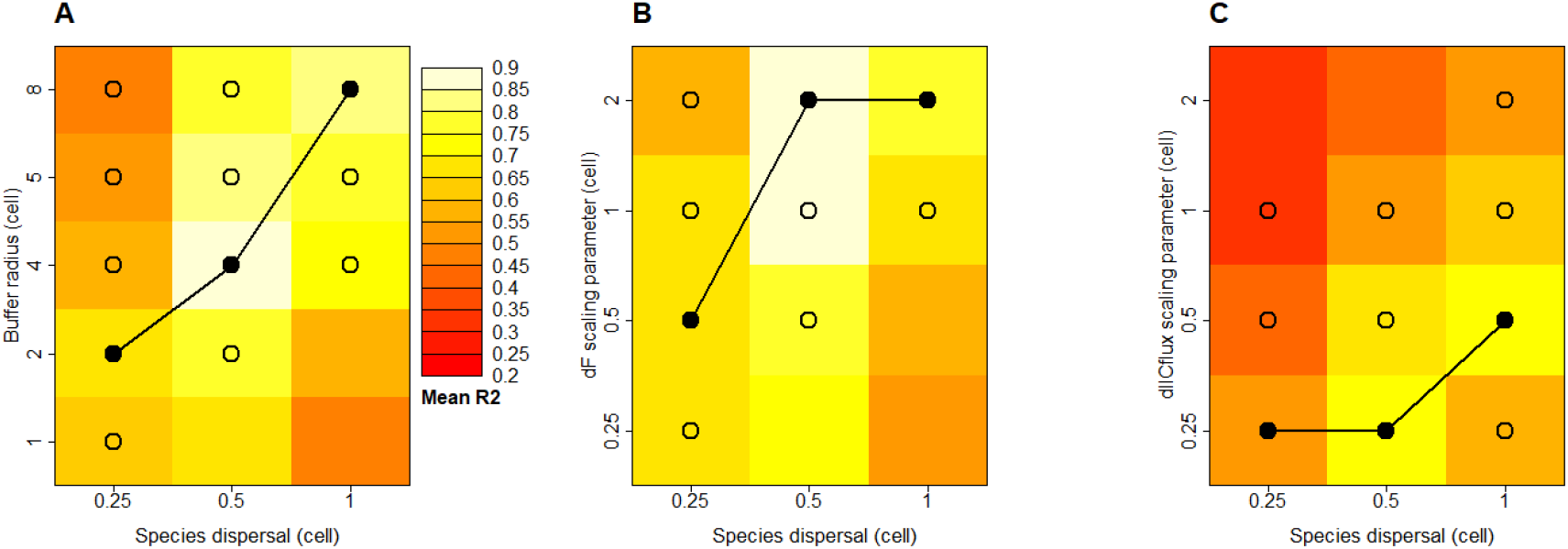
Combined effects of species dispersal distance and scaling parameter of patch connectivity indices on indices explanatory power. Panels A, B and C correspond to *Buffer*, *dF* and *dIICflux* indices respectively. Colors indicate the average explanatory power (*R2_spec_*) of the considered connectivity index across all the simulations with given species dispersal distance (*λ_s_*; x-axis) and scaling parameter (*r*_*buf*_ in panel A *λ*_*c*_ in panels B and C; y-axis). For each species dispersal distance, we marked with a black dot the “optimal” scaling parameter value, i.e. the scaling parameter value yielding the highest *R2*_*spec*_ among the explored values. We connected these dots to show, for each type of connectivity index, the species dispersal distance – scaling parameter relationship maximizing *R2*_*spec*_ in our simulations (beware that scales of axes are not linear). Because of our high number of simulations, the mean *R2*_*spec*_ obtained with optimal scaling is always significantly higher than mean *R2*_*spec*_ obtained with other scaling values. However, the difference between mean *R2*_*spec*_ for different scaling values was often small: for each species dispersal distance, we marked with circles the scaling parameter values that yield *R2*_*spec*_ values such that mean plus one standard deviation is higher than the mean *R2*_*spec*_ obtained with optimal scaling

### Hypothesis 3: connectivity variability effect

The linear model *logit(avR2_spec_)~log(avCV)* always detected a significant positive relationship between the coefficient variation and the explanatory power of connectivity indices (Fig. 4), with *p* < 2e-16 for *Buffer*, and *p* = 2e-9 and 4e-4 for *dF* and *dIICflux* respectively. The effect size of the coefficient of variation was markedly stronger for *Buffer* (estimate of 2.1 in the linear model with 0.17 s.d.) than for *dF* (estimate of 0.6 with 0.10 s.d.) and *dIICflux* (estimate of 0.3 with 0.09 s.d.). The *R^2^* of the linear model *logit(avR2_spec_)~log(avCV)* was stronger for *Buffer* (0.34) than for *dF* (0.13) and *dIICflux* (0.04), suggesting that the explanatory power of *Buffer* index is more tightly linked to its coefficient of variation than the two other indices. In line with this finding, adding the interaction of habitat aggregation and habitat proportion in the linear model (i.e. considering *logit(avR2_spec_)~log(avCV)+ Hurst exponent × habitat proportion*) did not significantly improve the fit for *Buffer* index (*p*=0.2 on a *F*-test),. By contrast, it did for the two other indices (*p*=2e-2 and *p*=4e-7 on a F-test for *dF* and *dIICflux* respectively). Note that, reciprocally, adding the coefficient of variation *avCV* always significantly improved the fit compared to a model with *Hurst exponent × habitat proportion* only (*p*<2e-16, *p*=2e-10 and *p*=5e-8 on a *F*-test for *Buffer*, *dF* and *dIICflux* respectively)

**Figure 4.**
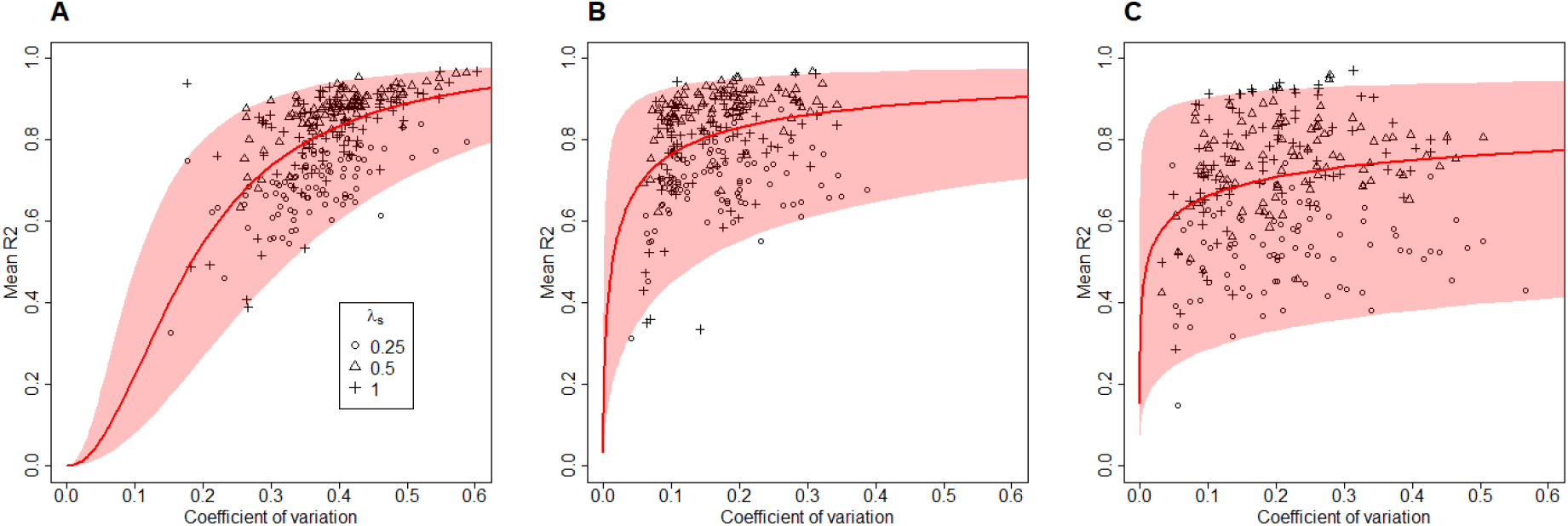
Explanatory power of patch connectivity indices (R2_spec_) as a function of the coefficient of variation of patch connectivity index. Panels A, B and C correspond to *Buffer*, *dF* and *dIICflux* index respectively. Symbols corresponds to species dispersal distance. The y-coordinate presents the average value of *R2*_*spec*_ with optimal scaling across the 10 community replicates associated to one landscape and one dispersal distance level. The x-coordinate presents the corresponding average coefficient of variation (CV) of the patch connectivity index with optimal scaling. Thus there are 3 Hurst exponents × 3 habitat proportions × 3 dispersal distances × 10 landscape replicates = 270 dots in each panel. The red curve present the fit of the linear model *logit(R2spec)~log(CV)* over these dots. The light-red envelope present a 95%-confidence interval around the fit

### Global performance of indices

When considering only connectivity indices with optimal scaling at fine patch delineation, a 95% of the 8,100 corresponding *R2*_*spec*_ values ranged between 0.35 (2.5% percentile) and 0.96 (97.5% percentile), with median value of 0.79. *Buffer* and *dF* stood out as the most performant index on average. The average *R2*_*spec*_ of *Buffer* was *R2_spec_*=0.79 (s.e.=0.003). Average *R2*_*spec*_ for *dF* differed from *Buffer* by −0.01, which was not significant. By contrast, the average *R2*_*spec*_ for *dIICflux* index significantly differed from *Buffer* by −0.11 (*t*-test; *t*=−27, *p*<2e-16)

## Discussion

Our study aimed at clarifying how patch delineation procedure, scaling of connectivity indices and connectivity variability among habitat patches could affect the explanatory power of three patch connectivity indices on species richness within patches. Our goal was to identify methods and landscape contexts that would foster strong patch connectivity – species richness relationships and thus provides relevant tests of the TIB framework. We expected that a virtual study would offer favorable settings to monitor the effect of patch connectivity on patch species richness in that they only modelled dispersal processes (combined with demographic stochasticity), and would therefore maximize our ability to study how methodological choices and landscape features affect the explanatory power of patch structural connectivity (*R2_spec_*). This expectation was verified in our results: *R2*_*spec*_ was generally high but showed marked contrast among simulations, hence allowing us to identify clear patterns when testing our three main hypotheses

### Patch delineation should match community size

We expected the predictive power of connectivity indices to be optimal when patch delineation matches the geographical scale of studied communities, hence ensuring no within-patch dispersal limitation. To test this hypothesis, we compared *R2*_*spec*_ when considering each elementary cell as a patch (the appropriate delineation with respect to simulations) versus when considering sets of contiguous cells as patches. *R2*_*spec*_ values were higher at fine patch delineation, where no dispersal limitation occurred within patches. The coarser patch delineation considering sets of contiguous habitat as patches led to important drop of *R2*_*spec*_ values, reaching about −0.2 when species harbored strong dispersal limitation (Fig. 2B). Our hypothesis was therefore corroborated

In the light of our results, we suggest that even when target habitats form “intuitive” patches (e.g. forest patches in agricultural landscapes), one should define *a priori* a grid with appropriate mesh size and use it to decompose the habitat map in elementary units, used for both community sampling and computation of connectivity indices. In particular, we discourage comparing community sampling and patch connectivity obtained at different spatial resolutions, which is often the case in empirical studies where species richness is derived from sampling covering only a small fraction of large patches obtained from coarse delineation (e.g. vegetation quadrats, birds point counts or insect traps). Our results suggest that using mesh size equal to the scale of dispersal distance for target organisms should allow strong patch connectivity - species richness relationships

Whether using even smaller mesh size should erode species richness – connectivity relationship remains an open question. It is however obvious that very fine patch delineation can make connectivity indices computation challenging, since it can increase by several orders of magnitude the number of spatial units. This particularly affects indices stemming from graph theory that need to determine shortest paths between all pairs of spatial units. Here we have been able to compute *dIICflux* in all the virtual landscapes at fine patch delineation (up to 4000 habitat units in a single landscape). Consequently, indices based on binary networks seem to pass the test of computational time. By contrast, we were unable to compute analogous indices in weighted networks (e.g., *dPCflux*; [30])

Determining *a priori* the appropriate mesh size corresponding to the scale of dispersal distance for the group of species under study is not straightforward, especially since in real communities – contrary to our simulations – movement capacity and dispersal distance are heterogeneous among species. Beyond the binary comparison between coarse and fine patch delineation that we proposed here, one should now explore the sensitivity of patch connectivity indices explanatory power to varying mesh size (as suggested by Mazerolle and Villard [29] in real empirical studies). This would allow assessing whether some degree of uncertainty on that parameter is acceptable

### Connectivity indices scaling should be adapted to species dispersal distance

Our second hypothesis was that the scaling of patch connectivity indices maximizing the explanatory power upon species richness should increase with dispersal distance of target organisms. We found that the scaling of patch connectivity indices leading to maximal explanatory power on species richness (the “scale of effect” *sensu* Jackson and Fahrig [19]) increased with the dispersal distance of target organism, in line with our hypothesis, and previous findings obtained from virtual studies [19,20]. This is a strong argument to prefer patch connectivity indices with a scaling parameter that can be modulated to match the dispersal distance of organisms rather than indices that cannot be adapted, like distance to nearest patch. It also confirms that the scale of effect should capture some quantitative features of species dispersal distance, as it is often contended in the empirical literature (e.g., [31,32])

However, the scale of effect should not be used as a quantitative estimate of dispersal distance for two reasons. First, we observed that scaling parameter values around the optimal one often generated a small drop of explanatory power, suggesting that the explanatory power was not highly sensitive to errors on scaling parameter value. Therefore, finding the scaling parameter that maximizes the correlation is probably not an accurate method to obtain estimate of species dispersal distance. This is consistent with the fact that, in empirical systems, buffer radii maximizing the explanatory power over species presence or abundance can spread over a large array of distances without significant drop of explanatory power, sometimes covering several orders of magnitude (e.g., [33]). Second, the quantitative relationship between the scale of effect and species dispersal distance was variable among indices tested. Rather, the relationship between the scale of effect and the scale of species dispersal distance may be used to roughly rank species or groups of species according to their dispersal distance. It can also contribute, when some *a priori* information is available about the dispersal distance of target organisms, to defining the range of scaling parameter values in which the scale of effect should be searched for Here we considered neutral metacommunities where all the species have the same dispersal distance. This greatly simplified the analysis of the relationship between the scale of effect of indices and species dispersal distances. However, species dispersal distances in real communities are known to be heterogeneous [34,35]. One may therefore question how our findings can transfer to real empirical studies. The fact that, for a given species dispersal distance and a given index, a broad range of scaling parameters have an explanatory power similar to the scale of effect could here turn out to be an advantage: a scaling parameter value adapted to the average dispersal distance of species in the community might be fairly adapted to all the species in the community. Of course, this should not be valid anymore if species dispersal distances are highly heterogeneous among species

### Buffer with appropriate scaling should be highly variable among patches

Our third and last hypothesis was that a higher variability of patch connectivity indices among sampled sites should increase the explanatory power of connectivity metrics upon species. We found a consistent positive relationship between the coefficient of variation of the patch connectivity indices and explanatory power, hence corroborating our expectation. The coefficient of variation of *Buffer* was sufficient to explain the fluctuation of explanatory power among landscapes with distinct habitat proportion or aggregation. Hence, the coefficient of variation of *Buffer* index with optimal scaling provides a remarkably simple and practical tool to assess whether a landscape has potential to reveal an effect of connectivity on species richness. Importantly, the relationship between the coefficient of variation and the explanatory power was looser for the two other index types explored (*dF* and *dIICflux*), and habitat aggregation and proportion seemed to affect the explanatory power beyond their effect on those indices’ variability. Thus, *Buffer* stands out as the appropriate index to assess connectivity variability in the context of our study. Whether this specificity holds when (i) broader range of scaling parameter values are explored for topological indices like *dIICflux* or (ii) landscapes harbor heterogeneous resistance is an open question that calls for more virtual studies. From an empirical perspective, our results emphasized the pivotal role of *Buffer* coefficient of variation and now call for defining appropriate thresholds on this coefficient to observe an effect on species richness. This requires meta-analyses of formerly published studies, accounting for taxa and habitat specificities

### Buffer and dF outcompete dIICflux in our simulations

Our study allowed us to compare the explanatory power of the three type of connectivity indices. *Buffer* and *dF* indices lead to high and very similar performance when used with appropriate scaling. This stemmed from the fact that these two indices are highly correlated (average correlation across landscapes above 0.95; Fig. S4). In our study, *Buffer* resembled *dF* index when its radius was about 4 times the *dF* scaling parameter value. [17] had already evidenced that correlations between IFM index (a generalization of the dF index; [38]) and buffers could reach 0.9 in a real landscape (their study did not focus on how the scaling of both indices could affect the correlation). *Buffer* and *dF* indices are both weighted sums of surrounding habitat cells contribution, where weights decreases with Euclidean distance following some kernel function. The only difference between the two indices is that *Buffer* is based on a step function while *dF* is based on a smoothly decreasing exponential kernel. We therefore interpret our results as the fact that changing the decreasing function used as a kernel may little affect the local connectivity as long as scaling is adjusted. This may explain why Miguet et al. [39] found that: (i) switching from buffer to continuously decreasing kernel little affected AIC or pseudo-R^2^ of models used to predict species abundances; (ii) neither continuously decreasing nor step function was uniformly better to explain species abundance across four case studies; (iii) different continuous shapes of kernel had quite indiscernible predictive performance

The *dIICflux* index had a lower explanatory power than *Buffer* and *dF* indices on average (−0.12 on *R_spec_*). This difference in global performance was made possible by the fact that *dIICflux* harbored a different profile than *dF* and *Buffer* in landscapes (Fig. S4), because it considers topological rather than Euclidean distance to compute connectivity. The use of five scaling values only in our analysis calls for some caution in the interpretation of the *dIICflux* lower explanatory power. The optimal scaling value of *dIICflux* for low and intermediate dispersal distances seemed to lie below the lower limit of the range explored in our study. Consequently, the explanatory power of this index might be underestimated compared to the other ones and partly explain why it seems less efficient in predicting species richness

Part of the relative success of *dF* and *Buffer* over *dIICflux* may also stem from the fact that we did not include different resistance values to habitat and matrix cells. When heterogeneous resistance occurs, landscape connectivity including displacement costs (e.g. least cost path, circuit theory) can be markedly different from connectivity based on Euclidean distance only [40], and may better capture the movement of organisms in real case study [41,42]. This probably also applies to patch connectivity. By connecting only cells that contain habitat, *dIICflux* and other indices based on topological distance within a graph could prove more performant when matrix has high resistance cost, and we may not find the same superiority of Euclidean indices as in our simulations

### Conclusion

Our results suggest that finding a strong effect of some patch structural connectivity on local species richness can occur only if: (i) spatial units used as patches are sufficiently small to prevent internal dispersal limitation within patches, which can be obtained by using a raster perspective with appropriate mesh size for patch delineation; (ii) the scaling of the patch connectivity index is adapted to the dispersal distance of species considered, which can be obtained by screening scaling parameters over a range of values defined from a priori knowledge about species dispersal distance; (iii) a *Buffer* index with optimal scaling harbors a high variability among sampled patches. When those three criteria are met, the absence of effect of connectivity on species richness should be interpreted as contradicting TIB hypotheses. *Buffer* indices particularly stood out in our analysis, as they efficiently summarized landscape effects on species richness and show higher explanatory power than other index types. When used with appropriate scaling, they seem a robust choice to recommend for empirical applications. However, new virtual studies including heterogeneous resistance within landscapes are necessary to ascertain this point

## Data accessibility

Virtual datasets combining simulation outputs and patch connectivity indices presented above have been made available on an online repository (doi: 10.5281/zenodo.3756712)

## Supplementary material

Codes for landscape generation and metacommunity simulation and codes of the analyses of virtual datasets presented above have been made available on an online repository (doi: 10.5281/zenodo.3756712)

## Acknowledgements

We thank Santiago Saura for sharing CONEFOR code. This work was part of the Patrames project funded by the convention 2016 n°2101870336 of the French Ministry of Environment (Water and Biodiversity Direction). Version 5 of this preprint has been peer-reviewed and recommended by Peer Community In Ecology (https://doi.org/10.24072/pci.ecology.100055)

## Appendix

**Figure S1.**
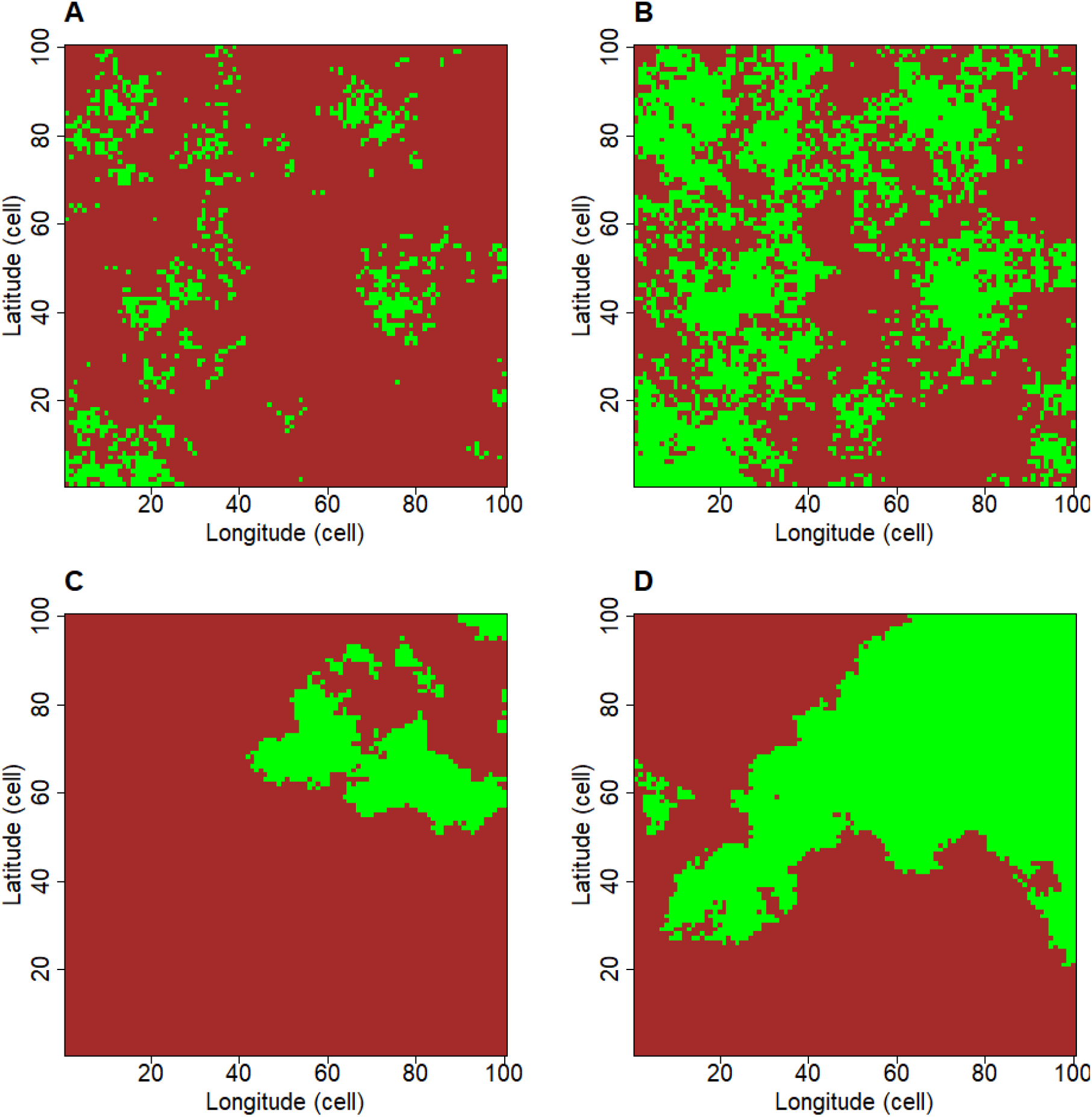
Simulated landscapes with contrasted aggregation and habitat proportion obtained from the midpoint displacement algorithm. Habitat is pictured in green, matrix in brown. Columns correspond to distinct levels of habitat proportion: panels A and C correspond to low habitat proportion (10%), panels B and D correspond to high habitat proportion (40%). Lines correspond to distinct habitat aggregation: panels A and B correspond to low habitat aggregation (Hurst exponent = 0.1), panels C and D correspond to high habitat aggregation (Hurst exponent = 0.9)

**Figure S2.**
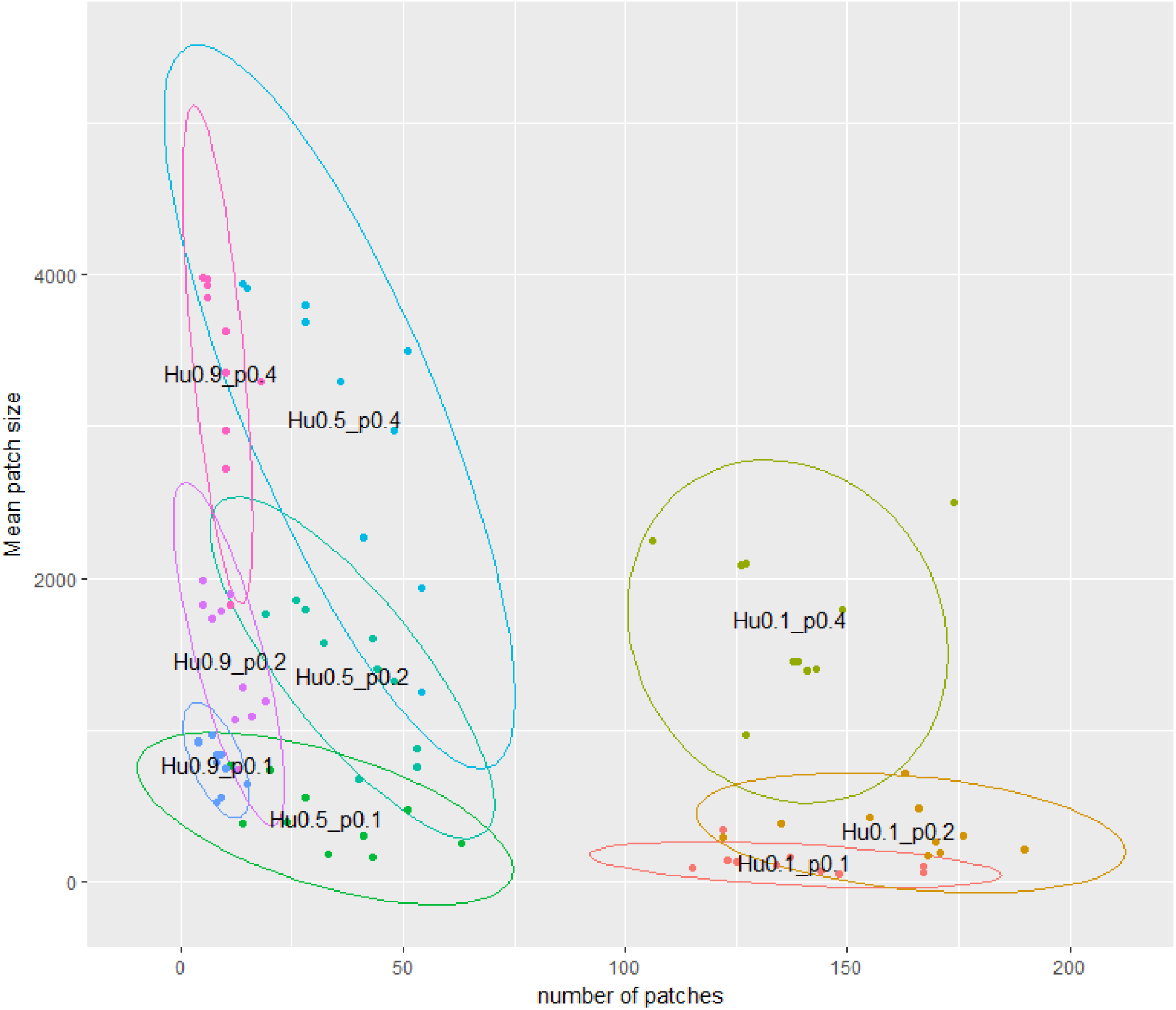
Average size and number of sets of contiguous cells within simulated landscapes. Colors correspond to distinct combinations of Hurst exponent (“Hu” in the labels) and habitat proportion (“p” in the labels). Ellipses correspond to 95%-CI of a fitted bivariate Student distribution

**Figure S3.**
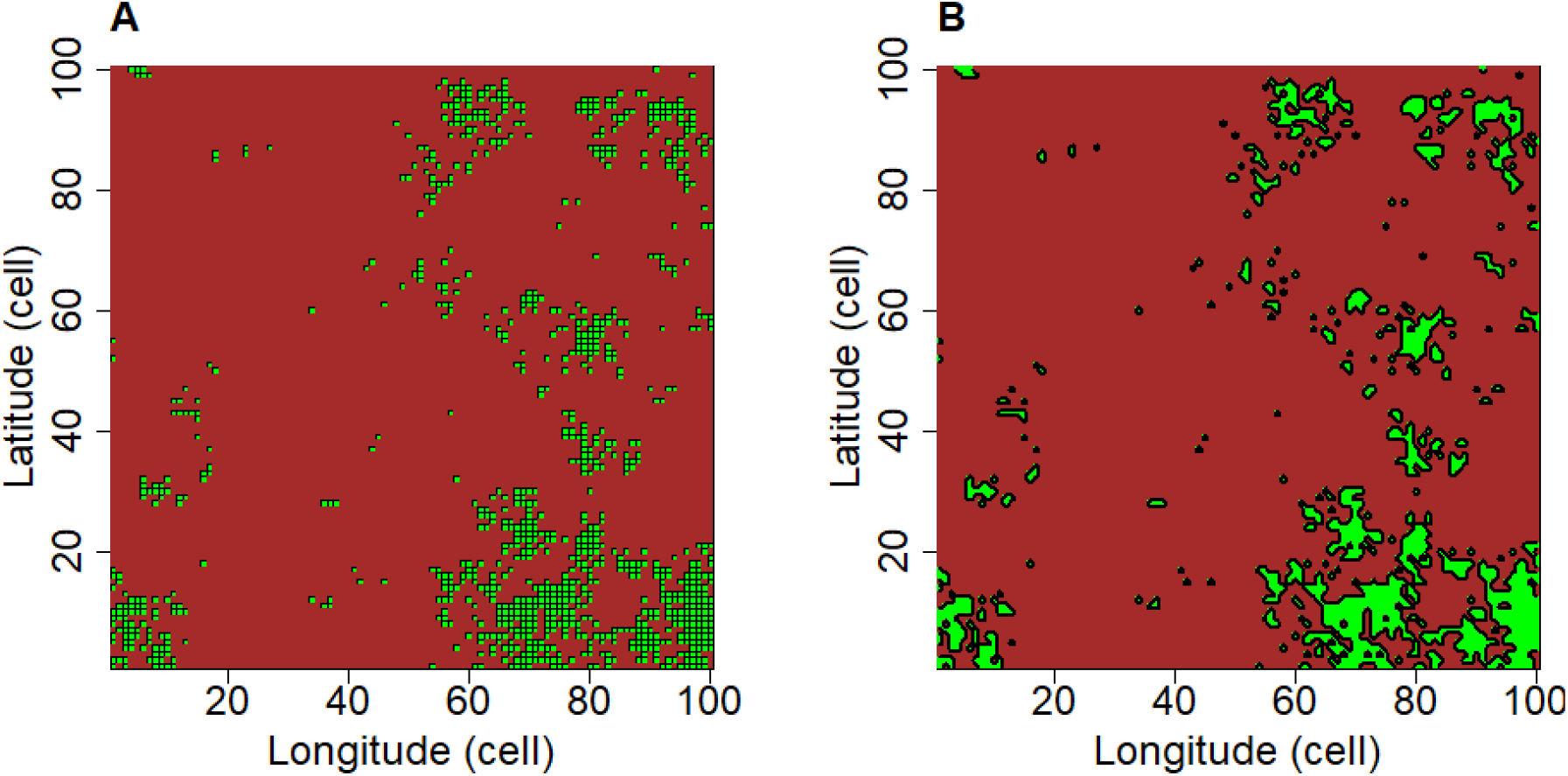
Lumping of contiguous cells generating the coarse patch delineation perspective. Panel A shows a habitat map where fine delineation of patches has been applied. Panel B shows the same habitat map where coarse patch delineation has been applied, i.e. sets of contiguous cells has been lumped together. Contiguity is based on the Von Neuman neighborhood of cells

**Figure S4.**
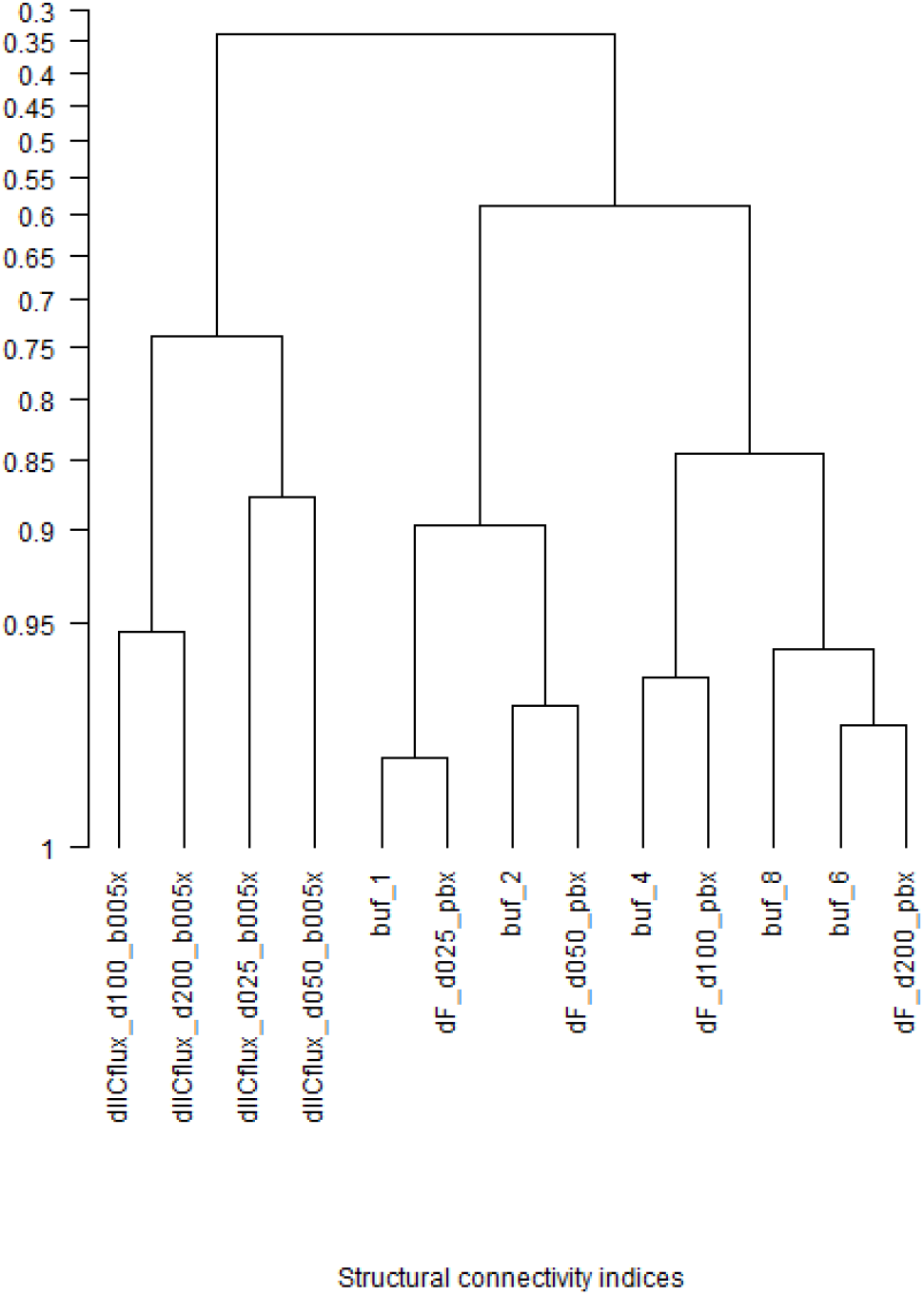
Dendrogram of Pearson correlation coefficients among patch structural connectivity indices across all landscapes. We presented correlations among *Buffer*, *dF* and *dIICflux* using ascending hierarchical classification. Within each of the 90 simulated landscapes, we computed the values of the 13 indices (accounting for distinct scaling values) in all habitat cells, which yielded 13 vectors of length 1000 to 4000 depending on the habitat proportion. We scaled each of the 13 vectors to mean 0 and variance 1, divided them by the square root of the number of habitat cells in the landscapes and computed pairwise Euclidean distances among them. We thus obtained one 13×13 distance matrix among patch connectivity indices in each of the 90 landscapes. Note that the distance between two indices corresponds to 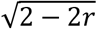, where *r* is the Pearson correlation between the indices across all habitat cells of the considered landscapes. We then averaged the 90 distance matrices to obtain one single 13×13 distance matrix as a basis for classification. We ran an ascending non-supervised classification (*hclust* function of R *base* package), using the *complete* method for group merging. A monophyletic group G with common ancestor located at value r means that any pair of indices within G has a correlation above r. Indices labels in the dendrogram are made of three parts separated by underscores “_”. The first part of the name indicates the type of the index (“buf”, “dF”, “dIICflux”). The second part of the name indicates the scale parameter of the index (“d025”, “d050”, “d100”, “d200” corresponding to *λ*_*c*_ = 0.25, 0.5, 1, 2 cells respectively, and “1”, “2”, “4”, “6”, “8” corresponding to *Buffer* radius *r_buf_* in cells). The last part in meaningless here

## References

1. MacArthur RH, Wilson EO. 1967 The theory of island biogeography. Princeton University Press. Princeton, NJ

2. Itescu Y. 2019 Are island-like systems biologically similar to islands? A review of the evidence. Ecography 42, 1298–1314. (doi:10.1111/ecog.03951)

3. Haila Y. 2002 A conceptual genealogy of fragmentation research: from island biogeography to landscape ecology. Ecological Applications 12, 321–334. (doi:10.1890/1051-0761(2002)012[0321:ACGOFR]2.0.CO;2)

4. Tischendorf L, Fahrig L. 2001 On the use of connectivity measures in spatial ecology. A reply. Oikos 95, 152–155. (doi:10.1034/j.1600-0706.2001.950117.x)

5. Schoener TW. 2010 The MacArthur-Wilson Equilibrium Model. In The theory of island biogeography revisited, pp. 52–87. Princeton, New Jersey: Princeton University Press

6. Jones NT, Germain RM, Grainger TN, Hall AM, Baldwin L, Gilbert B. 2015 Dispersal mode mediates the effect of patch size and patch connectivity on metacommunity diversity. Journal of Ecology 103, 935–944. (doi:10.1111/1365-2745.12405)

7. Löbel S, Snäll T, Rydin H. 2009 Mating system, reproduction mode and diaspore size affect metacommunity diversity. Journal of Ecology 97, 176–185. (doi:10.1111/j.1365-2745.2008.01459.x)

8. Mestre L, Jansson N, Ranius T. 2018 Saproxylic biodiversity and decomposition rate decrease with small-scale isolation of tree hollows. Biological Conservation 227, 226–232. (doi:10.1016/j.biocon.2018.09.023)

9. Buse J, Entling MH, Ranius T, Assmann T. 2016 Response of saproxylic beetles to small-scale habitat connectivity depends on trophic levels. Landscape Ecology 31, 939–949. (doi:10.1007/s10980-015-0309-y)

10. Schüepp C, Herrmann JD, Herzog F, Schmidt-Entling MH. 2011 Differential effects of habitat isolation and landscape composition on wasps, bees, and their enemies. Oecologia 165, 713–721. (doi:10.1007/s00442-010-1746-6)

11. Sverdrup-Thygeson A, Skarpaas O, Blumentrath S, Birkemoe T, Evju M. 2017 Habitat connectivity affects specialist species richness more than generalists in veteran trees. Forest Ecology and Management 403, 96–102. (doi:10.1016/j.foreco.2017.08.003)

12. Ranius T, Martikainen P, Kouki J. 2011 Colonisation of ephemeral forest habitats by specialised species: beetles and bugs associated with recently dead aspen wood. Biodiversity and Conservation 20, 2903–2915. (doi:10.1007/s10531-011-0124-y)

13. Alstad AO, Damschen EI. 2016 Fire may mediate effects of landscape connectivity on plant community richness in prairie remnants. Ecography 39, 36–42. (doi:10.1111/ecog.01492)

14. Prugh LR, Hodges KE, Sinclair ARE, Brashares JS. 2008 Effect of habitat area and isolation on fragmented animal populations. Proc Natl Acad Sci USA 105, 20770. (doi:10.1073/pnas.0806080105)

15. Thornton DH, Branch LC, Sunquist ME. 2011 The influence of landscape, patch, and within-patch factors on species presence and abundance: a review of focal patch studies. Landscape Ecology 26, 7–18. (doi:10.1007/s10980-010-9549-z)

16. Cook WM, Lane KT, Foster BL, Holt RD. 2002 Island theory, matrix effects and species richness patterns in habitat fragments. Ecology Letters 5, 619–623. (doi:10.1046/j.1461-0248.2002.00366.x)

17. Moilanen A, Nieminen M. 2002 Simple connectivity measures in spatial ecology. Ecology 83, 1131–1145. (doi:10.1890/0012-9658(2002)083[1131:SCMISE]2.0.CO;2)

18. Vieira MV, Almeida-Gomes M, Delciellos AC, Cerqueira R, Crouzeilles R. 2018 Fair tests of the habitat amount hypothesis require appropriate metrics of patch isolation: An example with small mammals in the Brazilian Atlantic Forest. Biological Conservation 226, 264–270. (doi:10.1016/j.biocon.2018.08.008)

19. Jackson HB, Fahrig L. 2012 What size is a biologically relevant landscape? Landscape Ecology 27, 929–941. (doi:10.1007/s10980-012-9757-9)

20. Economo EP, Keitt TH. 2010 Network isolation and local diversity in neutral metacommunities. Oikos 119, 1355–1363

21. Urban D, Keitt T. 2001 Landscape connectivity: a graph-theoretic perspective. Ecology 82, 1205–1218. (doi:10.1890/0012-9658(2001)082[1205:LCAGTP]2.0.CO;2)

22. Wiens JA, Stenseth NC, Van Horne B, Ims RA. 1993 Ecological Mechanisms and Landscape Ecology. Oikos 66, 369–380. (doi:10.2307/3544931)

23. Zurell D et al. 2010 The virtual ecologist approach: simulating data and observers. Oikos 119, 622–635. (doi:10.1111/j.1600-0706.2009.18284.x)

24. Hubbell SP. 2001 The unified neutral theory of biodiversity and biogeography. Princeton, NJ: Princeton University Press

25. Etherington TR, Holland EP, O’Sullivan D. 2015 NLMpy: a python software package for the creation of neutral landscape models within a general numerical framework. Methods in Ecology and Evolution 6, 164–168. (doi:10.1111/2041-210X.12308)

26. Bunn AG, Urban DL, Keitt TH. 2000 Landscape connectivity: A conservation application of graph theory. Journal of Environmental Management 59, 265–278. (doi:10.1006/jema.2000.0373)

27. Saura S, Torné J. 2009 Conefor Sensinode 2.2: A software package for quantifying the importance of habitat patches for landscape connectivity. Environmental Modelling & Software 24, 135–139. (doi:10.1016/j.envsoft.2008.05.005)

28. Saura S, Rubio L. 2010 A common currency for the different ways in which patches and links can contribute to habitat availability and connectivity in the landscape. Ecography 33, 523–537. (doi:10.1111/j.1600-0587.2009.05760.x)

29. Mazerolle MJ, Villard M-A. 1999 Patch characteristics and landscape context as predictors of species presence and abundance: A review1. Écoscience 6, 117–124. (doi:10.1080/11956860.1999.11952204)

30. Saura S, Pascual-Hortal L. 2007 A new habitat availability index to integrate connectivity in landscape conservation planning: Comparison with existing indices and application to a case study. Landscape and Urban Planning 83, 91–103. (doi:10.1016/j.landurbplan.2007.03.005)

31. Ranius T. 2006 Measuring the dispersal of saproxylic insects: a key characteristic for their conservation. Population Ecology 48, 177–188

32. Ranius T, Johansson V, Fahrig L. 2011 Predicting spatial occurrence of beetles and pseudoscorpions in hollow oaks in southeastern Sweden. Biodiversity and Conservation 20, 2027–2040. (doi:10.1007/s10531-011-0072-6)

33. Bergman K-O, Jansson N, Claesson K, Palmer MW, Milberg P. 2012 How much and at what scale? Multiscale analyses as decision support for conservation of saproxylic oak beetles. Forest Ecology and Management 265, 133–141. (doi:10.1016/j.foreco.2011.10.030)

34. Muller-Landau HC, Wright SJ, Calderón O, Condit R, Hubbell SP. 2008 Interspecific variation in primary seed dispersal in a tropical forest. Journal of Ecology 96, 653–667. (doi:10.1111/j.1365-2745.2008.01399.x)

35. Cadotte MW, Mai DV, Jantz S, Collins MD, Keele M, Drake JA. 2006 On Testing the Competition‐Colonization Trade‐Off in a Multispecies Assemblage. The American Naturalist 168, 704–709. (doi:10.1086/508296)

36. Calcagno V, Mouquet N, Jarne P, David P. 2006 Coexistence in a metacommunity: the competition– colonization trade‐off is not dead. Ecology Letters 9, 897–907

37. Laroche F, Jarne P, Perrot T, Massol F. 2016 The evolution of the competition–dispersal trade-off affects α- and β-diversity in a heterogeneous metacommunity. Proceedings of the Royal Society of London B: Biological Sciences 283. (doi:10.1098/rspb.2016.0548)

38. Hanski I. 1994 A practical model of metapopulation dynamics. Journal of Animal Ecology 63, 151–162. (doi:10.2307/5591)

39. Miguet P, Fahrig L, Lavigne C. 2017 How to quantify a distance-dependent landscape effect on a biological response. Methods in Ecology and Evolution 8, 1717–1724. (doi:10.1111/2041-210X.12830)

40. Simpkins CE, Dennis TE, Etherington TR, Perry GLW. 2018 Assessing the performance of common landscape connectivity metrics using a virtual ecologist approach. Ecological Modelling 367, 13–23. (doi:10.1016/j.ecolmodel.2017.11.001)

41. Emel SL, Storfer A. 2015 Landscape genetics and genetic structure of the southern torrent salamander, Rhyacotriton variegatus. Conservation Genetics 16, 209–221. (doi:10.1007/s10592-014-0653-5)

42. Vuilleumier S, Fontanillas P. 2007 Landscape structure affects dispersal in the greater white-toothed shrew: Inference between genetic and simulated ecological distances. Ecological Modelling 201, 369–376. (doi:10.1016/j.ecolmodel.2006.10.002)

43. Villard M-A, Metzger JP. 2014 REVIEW: Beyond the fragmentation debate: a conceptual model to predict when habitat configuration really matters. Journal of Applied Ecology 51, 309–318. (doi:10.1111/1365-2664.12190)

